# The relative effects of abiotic and biotic factors in explaining the structure of soil bacterial communities at diverse taxonomic levels

**DOI:** 10.1101/2024.10.29.620845

**Authors:** Baptiste Mayjonade, Rémy Zamar, Sébastien Carrere, Fabrice Roux

**Author notes:** **Competing Interests** The authors declare no competing financial interests.

## Abstract

Soil microbes play pivotal roles in the multifunctioning of terrestrial ecosystems. In the context of global changes, there is an urgent need to protect soil microbial diversity, which relies on determining the abiotic and biotic factors that influence the diversity, composition, and assemblage of soil microbiota. A large number of informative studies have reported edaphic properties and climate factors as key drivers of soil bacteria microbiota. However, these studies were mainly conducted at the phylum level and based on a restricted number of non-microbial variables. In this study, we aimed to estimate the relative effects of abiotic and biotic factors in shaping soil bacterial communities at diverse taxonomic levels by focusing on 160 natural sites located in the southwest of France for which a large and unique set of non-microbial variables is available. After characterizing soil bacterial communities with the highly taxonomically resolving *gyrB* gene, we identified that in addition to pH, temperature, and precipitations, soil bacterial communities at the lowest taxonomic levels appear strongly structured by soil micronutrients, notably manganese. On the other hand, soil bacterial communities at the highest taxonomic levels appear strongly structured by the interplay between descriptors of plant communities and edaphic properties. Similar to previous observations on microbial pathogens, the strong and positive associations between soil bacterial species and the presence of particular plant species suggest host specificity for soil commensal bacteria. Altogether, a deeper characterization of both abiotic and biotic factors could help fuel programs designed for protecting and restoring soil ecosystem functions.

## INTRODUCTION

Soil microbes play pivotal roles in the multifunctioning of terrestrial natural and managed ecosystems and contribute to a large number of key ecosystem processes, including nutrient cycling, carbon cycling and sequestration, soil structure and maintenance, water purification, degradation of pollutants, and ecosystem resilience to stresses [1–8]. These soil microbiota functions, in turn, confer either directly or indirectly a large number of benefits such as (i) climate regulation by consumption of carbon, methane, and nitrous oxide, three dominant gases responsible for 98% of the increased global warming [9–13], and (ii) regulation of the diversity and functioning of plant communities through, for instance, pathogen resistance, plant nutrition, and plant productivity [2, 14–16]. The soil microbiota–associated benefits ultimately contribute to animal and human health and well-being, which led to the proposal to include soil microbiota in the One Health policy [15, 17, 18].

There is therefore an urgent need to protect soil microbial diversity and to develop programs for personalized restoration of soil ecosystem functions [3, 9, 18], through for instance the use of inoculants as environmental probiotics or changes in land management practices [19–23]. Beyond the most precise possible taxonomic and/or functional description of the microbial diversity present in soils, protecting soil microbial diversity and developing restoration programs rely on determining the abiotic and biotic factors that influence the diversity, composition, and assemblage of soil microbiota [23, 24]. Identifying factors that regulate soil microbiota is even more relevant in the context of global changes that can directly threaten the contributions of soil microbes to ecosystem services and the health of aboveground biotic communities [12, 15, 19]. For instance, climate change can directly alter soil water regime and mineral composition evolution[25]. The increased occurrences of drought can reduce soil microbial diversity and abundance [26], with potential cascading effects such as an alteration in the rate of soil organic matter formation and degradation and a reduction in soil organic carbon content [27]. Also, temperature elevation together with altered humidity increases the proportion of soil-borne pathogens [28], and consequently the number and severity of epidemics [29]. In addition, the observed geographic range shifts of plant species under climate change can alter aboveground-belowground interactions [6, 30, 31].

Regarding bacterial communities, edaphic properties such as pH, soil organic carbon content, soil moisture, nitrogen availability, and redox status, emerge as the key drivers of soil bacterial microbiota, whatever the geographical scale considered [4, 5, 15, 16, 24, 32–38]. On the other hand, contrasting results have been observed on the relative importance of climate factors [26, 39–41], diversity and composition of plant communities [5, 6, 20, 31, 42, 43], and land management practices [20, 22, 36, 37] in driving the structure and function of soil bacterial communities. While informative, these studies, however, often rely on a restricted characterization of non-microbial factors [44]. For instance, despite the importance of micronutrients for soil fertility, only recent studies have reported the role of micronutrients in modulating the structure and function of soil bacterial communities [44–46]. Similarly, plant communities are generally classified into broad categories of vegetative type [1, 5, 6, 35], thereby impeding estimating whether particular plant species can be potential drivers of soil bacterial communities. In addition, the discrepancies observed among studies on the relative importance of climate factors, edaphic factors, and descriptors of plant communities in explaining soil bacterial communities may originate from a difference in the taxonomic scale at which the studies were conducted. Most studies estimated relationships between non-microbial factors and soil bacterial communities either at the phylum level or at the deepest taxonomic level provided by the genetic marker used for microbiota characterization. In line, during the last decade, soil bacterial communities have been mainly characterized using a metabarcoding approach based on markers designed on the hypervariable regions of the 16S rRNA gene. This marker has however a lower taxonomic resolution than other markers such as the *gyrB* gene that allows the distinction of bacterial operational taxonomic units (OTUs) down to the species level, and even to the phylotype level for some species such as the bacterial pathogen *Pseudomonas syringae* [47–50]. In addition, the *gyrB* gene is almost always in a single copy, thereby limiting the overestimation of the abundance of taxa carrying multiple copies of *rrn* operons [51].

In this study, we aimed to estimate the relative effects of abiotic and biotic factors in shaping soil bacterial communities at diverse taxonomic levels by amplifying a fraction of the *gyrB* gene. We focused on 160 natural sites inhabited by the model plant species *Arabidopsis thaliana* and located in the southwest of France (Fig. 1a) [49, 52]. These populations were chosen to maximize the diversity of habitats (such as climate, soil type, vegetation type, and degree of anthropogenic perturbation) encountered by *A. thaliana* [30, 49, 52]. For instance, despite the restricted size of the sampling area (∼8.2% of the total area of metropolitan France), this geographical region is under the influence of three contrasted climates (i.e., oceanic climate, Mediterranean climate, and mountain climate), with climate variation between the natural sites representing 24% of the climate variation observed among 521 European locations (from Spain to Finland) inhabited by *A. thaliana* [52]. In addition, the 160 natural sites have been previously characterized for 14 soil physicochemical variables (including micronutrients) and 49 descriptors of plant communities (including the absolute abundance of the 44 most prevalent plant species) [30].

**Fig. 1.**
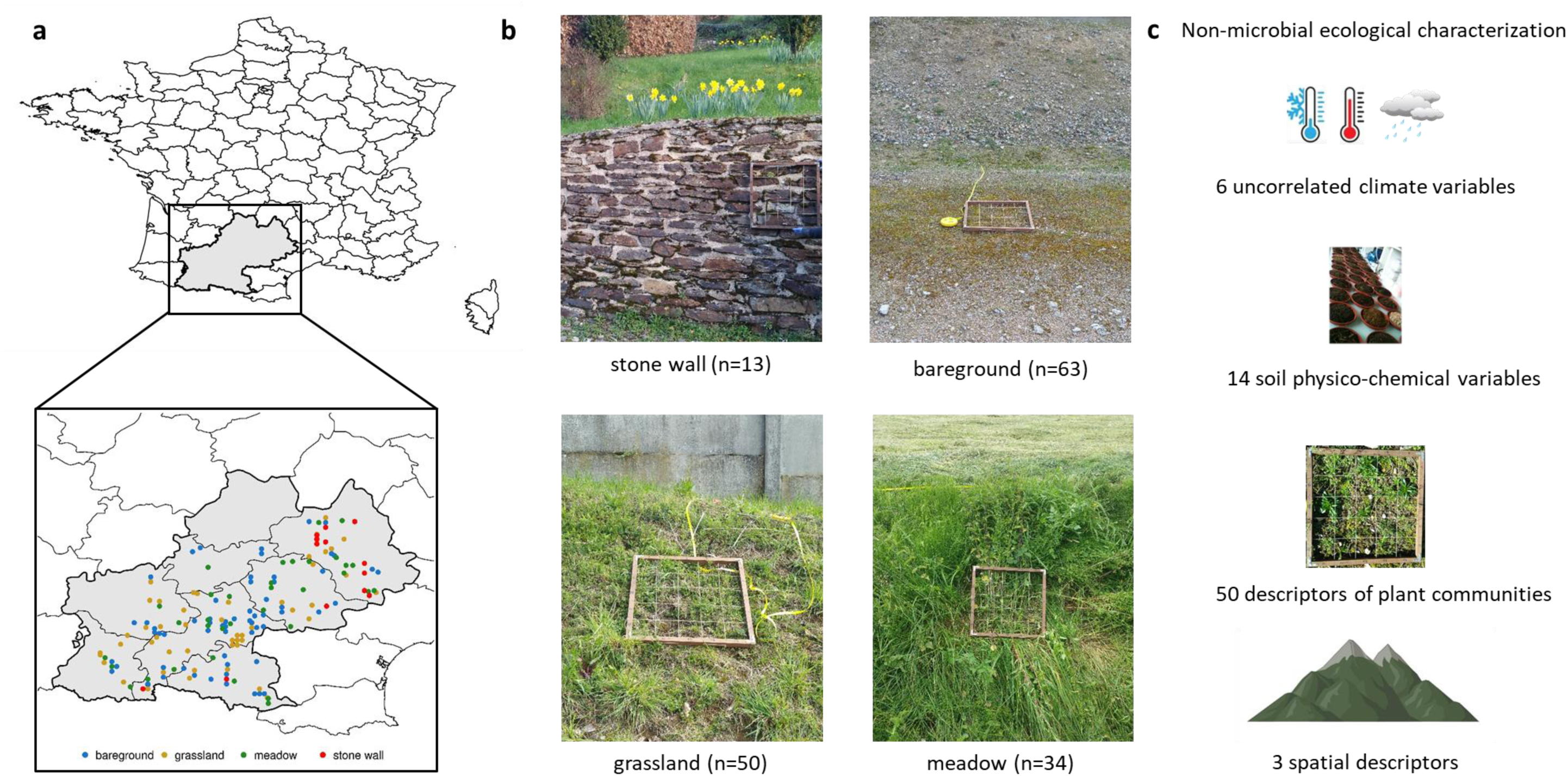
Location of the 160 sites inhabited by *Arabidopsis thaliana* in the southwest of France. **a** Mapping of the 160 sites according to four habitat categories. **b** Representative photos of the four habitat categories. The number of populations per habitat is indicated in brackets. **c** Non-microbial ecological characterization of the 160 sites, including 6 uncorrelated climate variables (two temperature-related variables and four precipitation-related variables), 14 soil physico-chemical variables describing the main soil agronomic properties related to plant growth, 50 descriptors of plant communities (richness, Shannon index, composition, plant cover, density of *A. thaliana*, presence/absence of the most prevalent companion plant species), and three spatial descriptors (latitude, longitude, and elevation).

## MATERIAL AND METHODS

### Study sites, ecological characterization, and soil sampling

In this study, we focused on 160 natural sites located in the southwest of France (Fig. 1a) inhabited by the model plant species *Arabidopsis thaliana* and classified into four habitat categories, i.e. stone wall, bare ground, grassland, and meadow (Fig. 1b, Supplementary Data 1) [30]. All these sites (except for the JACO-B site) have been previously characterized for a set of 6 uncorrelated climate variables (2003-2013 period) from the ClimateEU database, 14 soil physicochemical variables in 2015, 49 descriptors of plant communities in 2015, and 3 spatial descriptors [30, 49, 52] (Fig. 1c, Supplementary Data 1). In this study, we also considered plant cover that corresponds to the average percentage of surface covered by plants averaged between two 50 cm × 50 cm quadrats divided into 25 smaller squares (10 cm × 10 cm) [30] (Fig. 1c, Supplementary Data 1).

Because the soil environment is highly heterogeneous [23], collection and sampling of the soil compartment was performed following the experimental protocols described in [53, 54]. For each natural site, four soil cores (encompassing the 0-5 cm depth interval below the litter layer) were collected using flame-sterilized spoons and transferred on a sterilized ceramic plate. Soil samples were mixed by hand on the ceramic plate and sieved through a mesh of 5 mm using autoclaved plastic sieves. Gloves and the ceramic plate were sterilized by using Surface’SafeAnios®. Sieved soil samples were placed into sterilized Falcon 50mL tubes and immediately stored on ice. Back at the laboratory on the day of collection, three soil subsamples were individually stored in autoclaved 1.5mL Eppendorf tubes and stored at −20°C. For the purpose of a citizen project, a similar approach was adopted to collect soil samples from 56 common gardens located in the southwest of France (Supplementary Data 1). Soil sampling was performed in natural populations and common gardens in mid-November 2018.

### Soil microbial community DNA extraction and high-throughput sequencing

DNA was extracted for a total of 756 samples (160 natural sites × 3 subsamples + 56 common gardens × 4 subsamples + 13 common gardens × 4 additional subsamples). Total DNA was extracted using the FastDNA™ SPIN Kit for Soil (MP Biomedicals, Santa Ana, CA, USA) and following the manufacturer’s recommendations. For each DNA sample, the *gyrB* amplicon was amplified in a PCR reaction of 25 µl using the primers described in [47]. Each PCR reaction consisted of 2µl of DNA, 2.5µl of 10X MolTaq Buffer (Molzym GmbH & Co. KG, Bremen, Germany), 0.2µl of MolTaq 16S DNA polymerase, 0.5µl of 10mM dNTP (Promega, Madison, WI, USA), 0.25µl of 50mg/ml Non-Acetylated BSA (Thermofisher, Waltham, MA, USA), 0.75µl of 10µM HPSF forward, 0.75µl of 10µM HPSF reverse primer (Eurofins, Luxembourg) and 19.4µl of sterile biology molecular grade water. Cycling conditions consisted of an initial denaturation of 1 min at 95°C followed by 35 cycles of 95°C/30seconds, 55°C/1minute, 72°C/90seconds, and a final extension of 10 minutes at 72°C. Samples were randomized in eight 96-well PCR plates with other soil samples from another project. Each PCR reaction was repeated three times and pooled in a 96-well plate.

PCR products were purified with Agencourt®AMPure® magnetic beads following the manufacturer’s instructions. Purified amplicons were quantified with a NanoDrop 8000 and appropriately diluted to obtain an equimolar concentration. Two microliters of equimolar PCR-purified products were used for a second PCR with a universal P5 primer and 384 different P7 primers containing different indices used to multiplex samples. The second PCR amplicons were then purified and quantified as described above to obtain a unique equimolar pool. The latter was quantified by real-time quantitative PCR and then sequenced on a MiSeq PE250 platform (Illumina Inc., San Diego, CA, USA) in the GeT-PlaGe Platform (Toulouse, France). MS-102-3003 MiSeq Reagent Kit v3 600 cycle was used for this purpose.

### gyrase *β*-subunit gene amplicon data analysis

Raw sequences were trimmed to remove the first 20 5’ nucleotides corresponding to primer sequences and the last 3’ nucleotides, resulting in trimmed reads of 200 nucleotides. Amplicon Sequence Variants (ASVs) were obtained by running DADA2 software (version 1.18.0) [55] on these cleaned sequences. Taxonomic classification was assigned using a human-curated gyrB database composed of 38,929 sequences [47] (Supplementary Data 2). A final clustering step was performed using SWARM software (version 3.0.0) [56] to reduce redundancy. The taxonomic assignment of clusters was set to the most abundant ASV within the cluster.

We identified 327,387 ASVs across the 756 samples, including 241,184 ASVs for the 480 samples of the 160 natural sites (Supplementary Data 3). The ASV matrix related to natural sites being sparse, a Hellinger transformation [57] based on relative abundances was applied using the *vegan* R package version 2.5-7 prior β-diversity analyses. A permutational multivariate analysis of variance (PERMANOVA) performed on a Bray–Curtis dissimilarity matrix with the *adonis* function (*vegan* R package version 2.5-7) indicated that differences between the 160 natural sites explained 76.6% (*P* = 0.001) of bacterial community composition. This result indicated a strong repeatability between the three biological replicates per natural site. For each natural site, reads from the three biological replicates were therefore pooled for subsequent analyses.

### Statistical analysis

Shannon index for the 160 natural sites was estimated on the initial ASV matrix composed of 241,184 ASVs, by using the summary single function of mothur [58]. Differences in Shannon diversity between the four habitat categories were tested by an analysis of variance with the *aov* function under the *R* environment, followed by a Tukey’s HSD (Honestly Significant Difference) test. Spearman’s correlation coefficients were estimated between Shannon index and each of the 73 non-microbial ecological variables by using the *corr.test* function from the *psych* R package.

At each taxonomic level (i.e. phyla, class, order, family, genus, species, SWARM, and ASV), we estimated the relative abundance of each taxonomic group. After removing unclassified taxonomic groups, a Hellinger transformation [55] was performed on the matrix of relative abundances by using the *vegan* R package version 2.5-7. At each taxonomic level, differences in bacterial microbiota composition between the four habitat categories were tested by running a PERMANOVA on Bray–Curtis dissimilarity matrices with the *adonis* function (vegan R package version 2.5-7). At each taxonomic level, PERMANOVA was also used to test the effect of each of the 73 non-microbial ecological variables on bacterial microbiota composition.

At each taxonomic level, Spearman’s pairwise correlation coefficients were computed between each taxonomic group and the 73 non-microbial ecological variables by using the *corr.test* function from the psych R package. A correction for the number of tests was performed to control for the False Discovery Rate (FDR) at a nominal level of 5% by using the *p.adjust* function under the R environment. As a complementary approach to identifying non-microbial ecological variables associated with taxonomic groups, a sparse partial least-square regression (sPLSR) approach [59] was run on the same datasets by using the mixOmics package implemented in the *R* environment [60]. Following [49], the significance of the non-microbial ecological variables included in the linear combinations was estimated by a Jackknife resampling approach by leaving out 10% of the samples (i.e. natural populations of *A. thaliana*) 1000 times. Only non-microbial ecological variables with a loading value above 0.2 in more than 75% of the resampled matrices were considered significant. For estimating Spearman’s pairwise correlations and running sPLSR, only taxonomic groups present in more than 15 sites were considered.

## RESULTS

### Bacterial diversity, dominance, and prevalence

Based on the sequencing of a *gyrB* gene amplicon, we identified 27 phyla, 48 classes, 82 orders, 144 families, 372 genera, 563 species, 212,837 SWARMs, and 241,184 ASVs in the soil of 160 natural sites located in the southwest of France (Supplementary Data 3, Supplementary Data 4). The 14 phyla detected in more than 75% of the natural sites present a mean relative abundance per natural site ranging from 15.05% to 0.05% (Fig. 2). Betaproteobacteria, Alphaproteobacteria, Actinobacteria, Gammaproteobacteria, Bacteroidetes, Gemmatimonadetes and Acidobacteria were the most abundant (>3%) and prevalent (i.e. presence in all the natural sites) phyla (Fig. 2). In contrast, the rare phyla (e.g. Chlamydiae, Chloroflexi) were detected in fewer than 25% of the natural sites, with a mean relative abundance less than 0.01% per natural site (Fig.2).

**Fig. 2.**
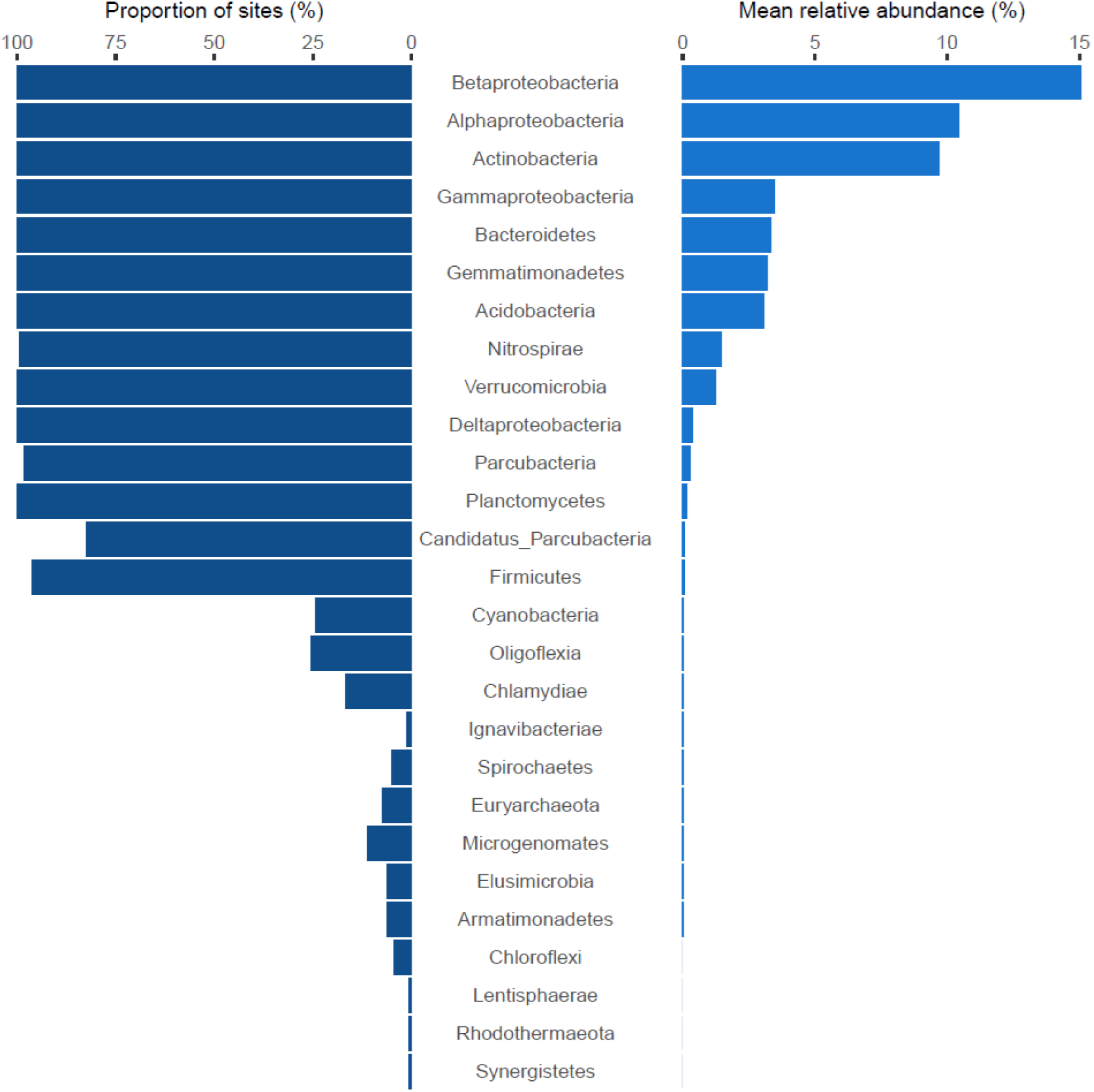
Representativeness of bacterial phyla in 160 soils from the southwest of France. Left: The proportion of sampling sites where phyla were present. Right: The mean relative abundance of the phyla.

### Relationships between bacterial alpha-diversity and environmental variables

Based on the 241,184 ASVs, we observed an extensive variation of bacterial alpha-diversity among the 160 natural sites, with Shannon index values ranging from 5.6 to 7.3 (Fig. 3a). Interestingly, Shannon index was significantly different among the four habitat categories (Fig. 3a). Specifically, Shannon index was significantly higher for soils collected in meadows than for soils collected on stone walls, with soils collected in bare ground and grassland habitats having intermediate values of Shannon index (Fig. 3a).

**Fig. 3.**
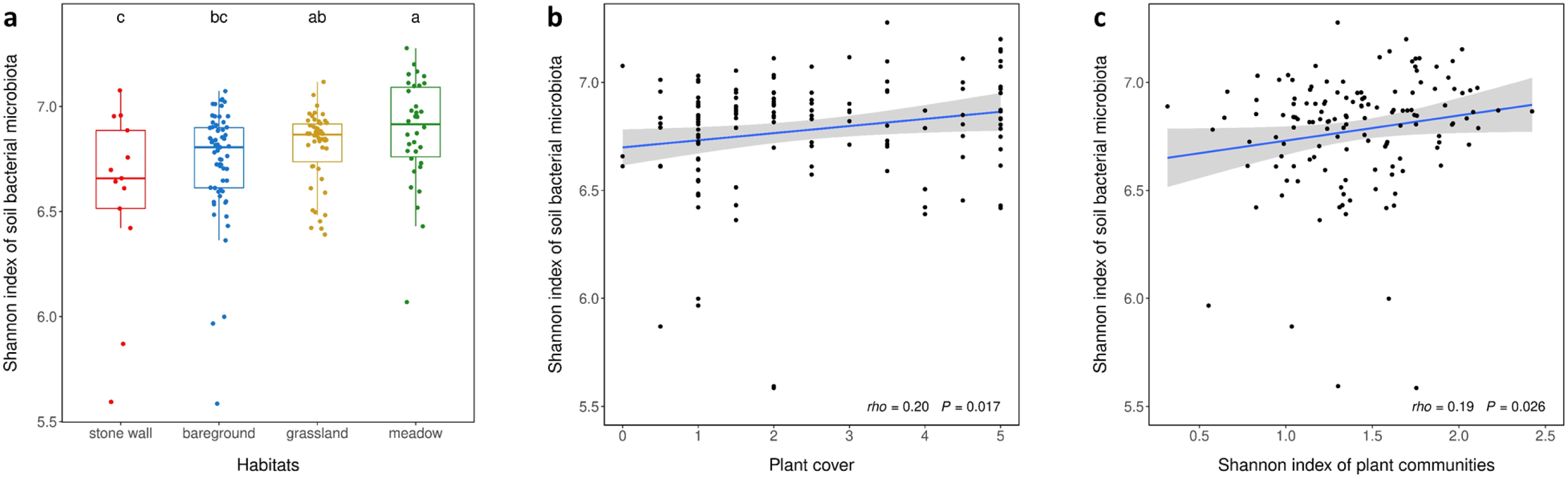
Association between Shannon index of soil bacterial communities and non-microbial ecological variables. **a** Effect of habitats on Shannon index of soil bacterial communities. Different upper letters indicate different groups after a Tukey HSD correction for multiple pairwise comparisons. **b** Positive relationship between Shannon index of soil bacterial communities and plant cover. **c** Positive relationship between Shannon index of soil bacterial communities and Shannon index of plant communities.

To identify non-microbial ecological variables underlying the difference in Shannon index among the four habitat categories, we tested the association of the Shannon index with 6 climate variables, 14 soil physicochemical properties, 50 descriptors of plant communities, and 3 spatial descriptors (Fig. 1c). While a positive association was detected between Shannon index of bacterial communities and Shannon index of plant communities (Fig. 3b), plant cover (Fig. 3c) and the presence of the plant species *Valerianella locusta* (Supplementary Figure S1), a negative association was detected between Shannon index of bacterial communities and the concentration in soil potassium (Supplementary Figure S1), the presence of the plant *Erophila verna* and the presence of a *Geranium* species (Supplementary Table S1).

### Relationships between bacterial composition and non-microbial environmental variables

At each taxonomic level, microbiota composition variance was weakly but significantly explained by differences among the four habitat categories (PERMANOVA, phylum: *R*² = 3.7%, *P* = 0.006; class: *R*² = 6.3%, *P* < 0.001; order: *R*² = 6.1%, *P* < 0.001; family: *R*² = 5.5%, *P* < 0.001; genus: *R*² = 4.5%, *P* < 0.001; species: *R*² = 4.4%, *P* < 0.001; SWARM: *R*² = 2.9%, *P* < 0.001; ASV: *R*² = 2.7%, *P* < 0.001). We then ran PERMANOVA to identify the main non-microbial ecological variables associated with microbiota composition variation. At each taxonomic level, most climate variables, soil physicochemical properties, and spatial descriptors were identified as explanatory variables but individually explained on average a small fraction (< 5%) of microbiota composition variance (Fig. 4, Supplementary Data 5). While very few descriptors of plant communities were associated with microbiota composition variation at the lowest taxonomic levels (i.e., from phylum to species), more than 20 descriptors of plant communities were identified as explanatory variables at the very highest taxonomic levels (i.e., SWARM and ASV) (Fig. 4, Supplementary Data 5). In addition, at the SWARM and ASV levels, descriptors of plant communities explained on average more microbiota composition variance than other non-microbial ecological variables, with some descriptors of plant communities individually explaining more than 15% of microbiota composition variance (Fig. 4, Supplementary Data 5). To further investigate the relationship between microbiota composition and non-microbial ecological factors, we then explored at each taxonomic level the relationship between the relative abundance of each microbial taxonomic group and each non-microbial ecological factor. For statistical power, we only considered taxonomic groups present in more than 15 sites, resulting in a final list of 18 phyla, 32 classes, 50 orders, 71 families, 130 genera, 127 species, 1010 SWARMs, and 769 ASVs (Supplementary Data 6). While our selection criteria led to a mean decrease of 62.2% in the number of taxonomic groups across the eight taxonomic levels, the remaining taxonomic groups represent on average 77.8% of the total number of reads (Supplementary Table S2). To identify relationships between microbial and non-microbial variables, we used two complementary approaches, that is, estimating Spearman’s pairwise correlations and running a sparse partial least-square regression (sPLSR) approach. While the first approach allows identifying non-linear relationships and is robust to outliers, the latter approach allows identifying linear combinations of non-microbial ecological factors associated with the variation of the relative abundance of a particular taxonomic group.

**Fig. 4.**
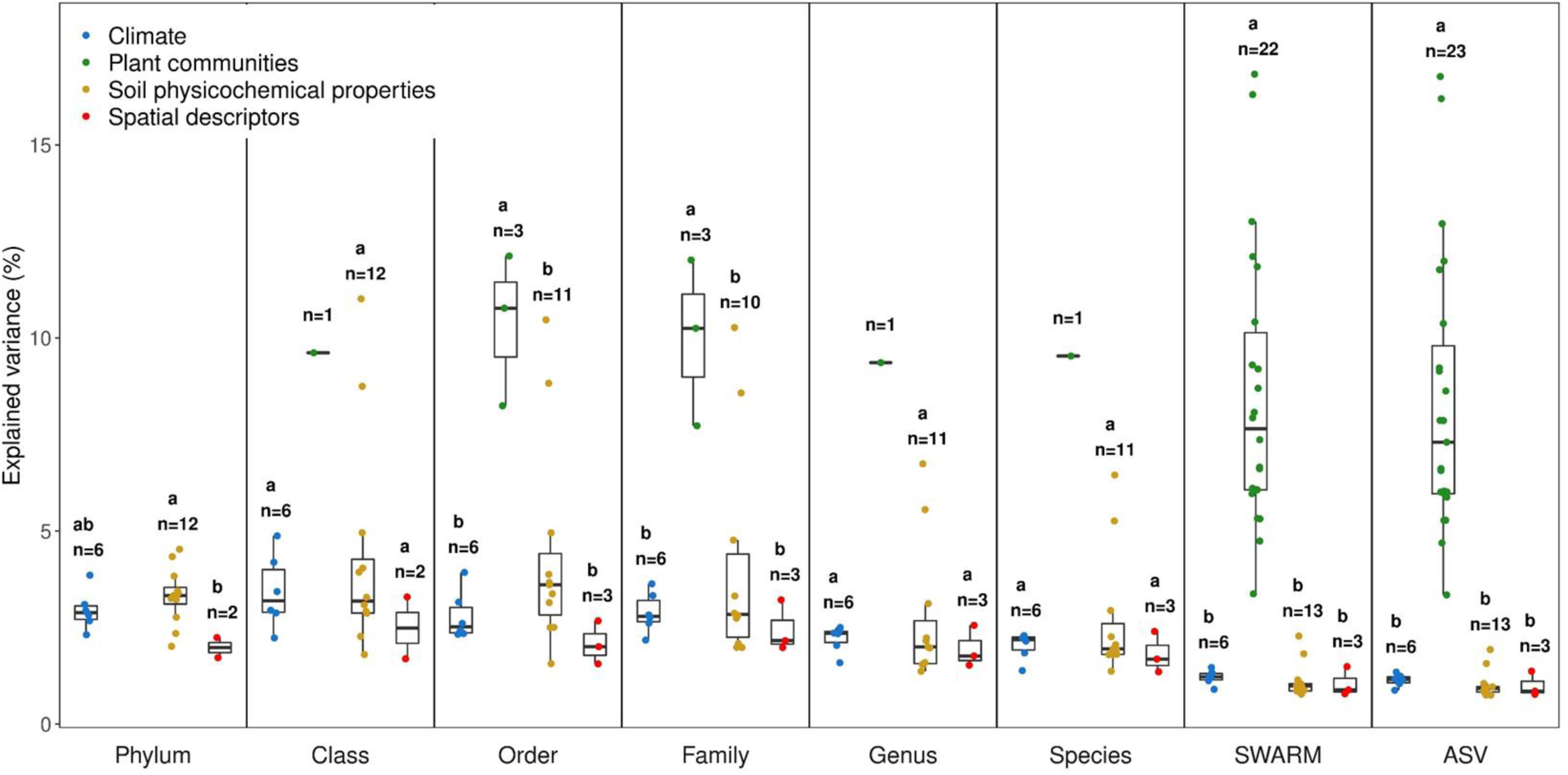
Identification of non-microbial ecological variables associated with microbiota composition variation at each of the eight taxonomic levels. Different upper letters indicate different groups after a Tukey HSD correction for multiple pairwise comparisons. Only non-microbial ecological categories with more than one non-microbial ecological variable were considered for running a Tukey HSD correction.

Both approaches revealed that the fraction of significant associations with individual microbial taxonomic groups decreased and increased from the lowest to the highest taxonomic levels for soil physicochemical properties and descriptors of plant communities, respectively (Fig. 5a, 5c, Supplementary Data 7, Supplementary Data 8). For instance, while the fraction of significant associations identified by sPLSR between individual microbial taxonomic groups and soil physicochemical properties decreased from 59% at the phylum level to 30.2% at the ASV level, the fraction of significant associations between individual microbial taxonomic groups and descriptors of plant communities increased from 19.7% at the phylum level to 48.7% at the ASV level (Fig. 5c, Supplementary Data 8). The fraction of significant associations between individual taxonomic groups and either climate variables or spatial descriptors remains stable across the eight taxonomic levels (Fig. 5a, 5c, Supplementary Data 7, Supplementary Data 8).

**Fig. 5.**
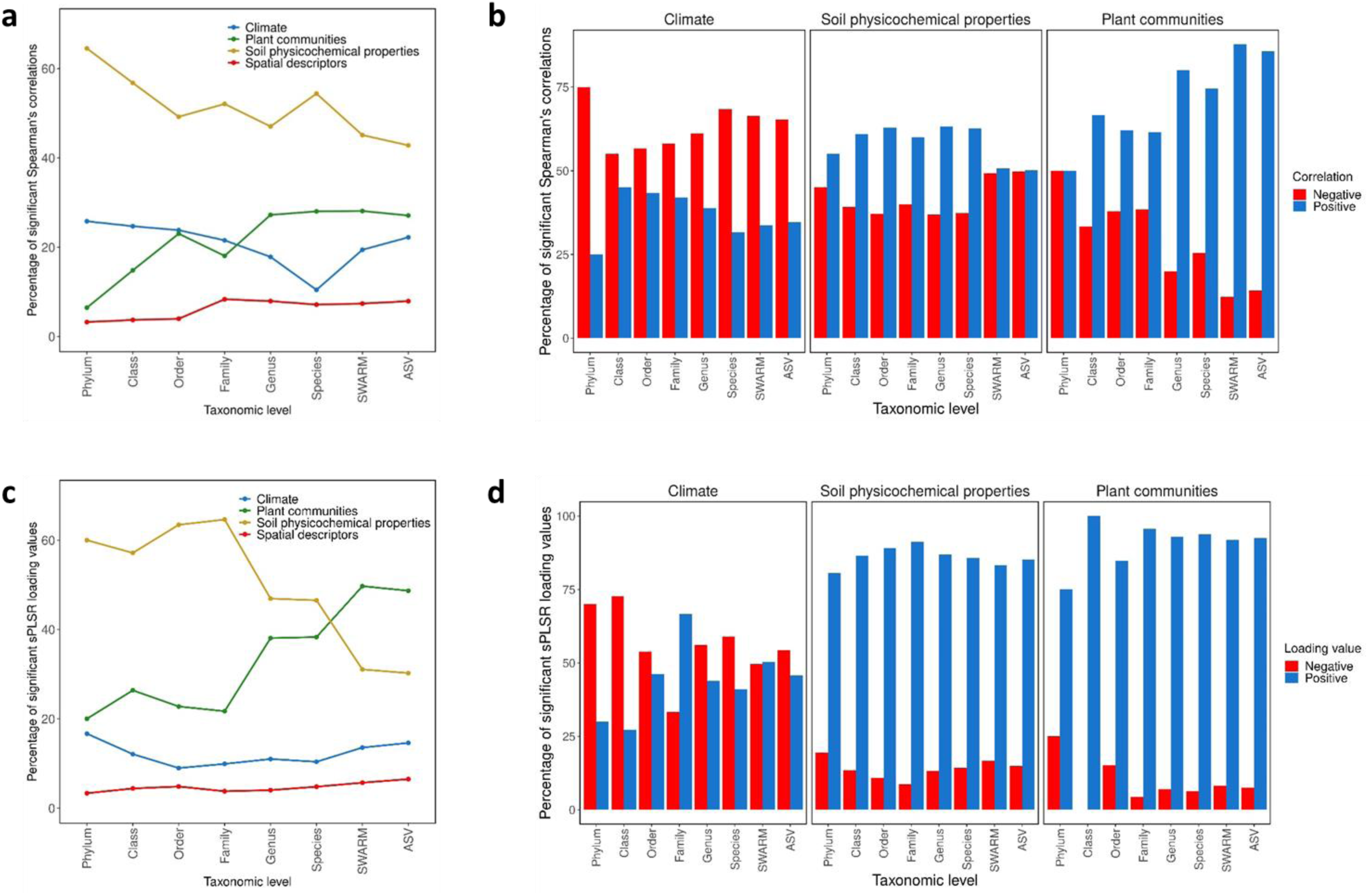
Identification of non-microbial ecological variables associated with individual taxonomic groups at each of the eight taxonomic levels. **a** Relative fraction between the four non-microbial ecological categories for the number of significant Spearman’s pairwise correlations computed between taxonomic groups and the 73 non-microbial ecological variables. **b** Relative fraction of positive and negative significant Spearman’s pairwise correlations at each taxonomic level and for the three non-microbial ecological categories ‘climate’, ‘soil physico-chemical properties’, and ‘plant communities’. **c** Relative fraction between the four non-microbial ecological categories for the number of non-microbial ecological variables with an absolute sPLSR loading value > 0.2. Only non-microbial ecological variables with a loading value above 0.2 in more than 75% of the 1000 Jackknife resampled matrices were considered significant. **d** Relative fraction of non-microbial ecological variables with a loading value > 0.2 (positive) or < −0.2 (negative), at each taxonomic level and for the three non-microbial ecological categories ‘climate’, ‘soil physico-chemical properties’, and ‘plant communities’.

While individual microbial taxonomic groups were more often negatively than positively associated with climate variables, individual microbial taxonomic groups were mainly positively associated with variation in soil physicochemical properties and descriptors of plant communities (Fig. 5b, 5d, Supplementary Data 7, Supplementary Data 8). In addition, the sPLSR approach revealed a higher fraction of positive associations between individual microbial taxonomic groups and either soil physicochemical properties or descriptors of plant communities than the approach based on estimating Spearman’s correlations (Fig. 5b, 5d). This suggests that individual microbial taxonomic groups are positively associated with linear combinations of soil physicochemical properties and descriptors of plant communities.

### Identification of key environmental variables associated with individual microbial taxonomic groups

Although mean annual temperature, spring precipitation, and fall precipitation appear as key climate variables associated with individual microbial taxonomic groups across the eight taxonomic levels, pH and soil magnesium concentration were much more often related to individual microbial taxonomic groups (Fig. 6). Another variable very often related to individual microbial taxonomic groups corresponds to elevation (Fig. 6). In addition, while the number of associations with individual microbial taxonomic groups was on average smaller for descriptors of plant communities than for climate variables and soil physicochemical properties (Fig. 6, Supplementary Data 7, Supplementary Data 8), the strength of association was significantly stronger for descriptors of plant communities than for climate variables, the strength of association being intermediate for soil physicochemical properties (Fig. 6, Supplementary Data 7, Supplementary Data 8). The diversity of associations between individual microbial taxonomic groups and non-microbial factors detected in this study is illustrated by nine ‘individual microbial taxonomic group × non-microbial ecological factor’ combinations (Fig. 7).

**Fig. 6.**
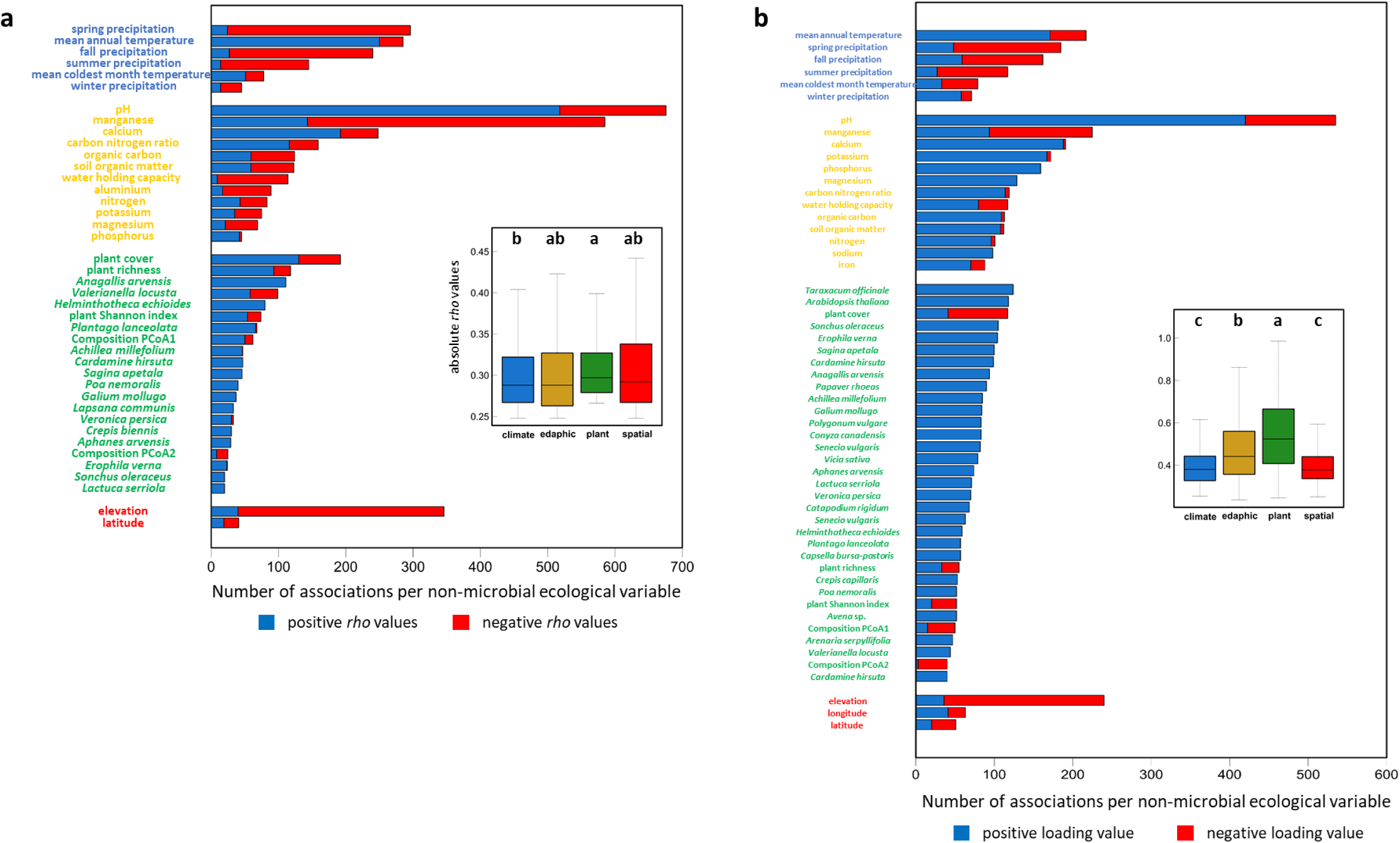
Number of taxonomic groups associated with each non-microbial ecological variable across the eight taxonomic levels. **a** Number of taxonomic groups with positive and negative Sperman’s correlation coefficients (*rho*) with each non-microbial ecological variable. Only non-microbial ecological variables associated with at least 20 taxonomic groups are depicted. Inner box plot: Variation in absolute *rho* values among the four categories of non-microbial ecological variables. Different letters indicate different groups according to the category after a Tukey correction for multiple pairwise comparisons. **b** Number of taxonomic groups with positive and negative loading values with each non-microbial ecological variable. Only non-microbial ecological variables associated with at least 40 taxonomic groups are depicted. Inner box plot: Variation in absolute loading values among the four categories of non-microbial ecological variables. Different letters indicate different groups according to the category after a Tukey correction for multiple pairwise comparisons.

**Fig. 7.**
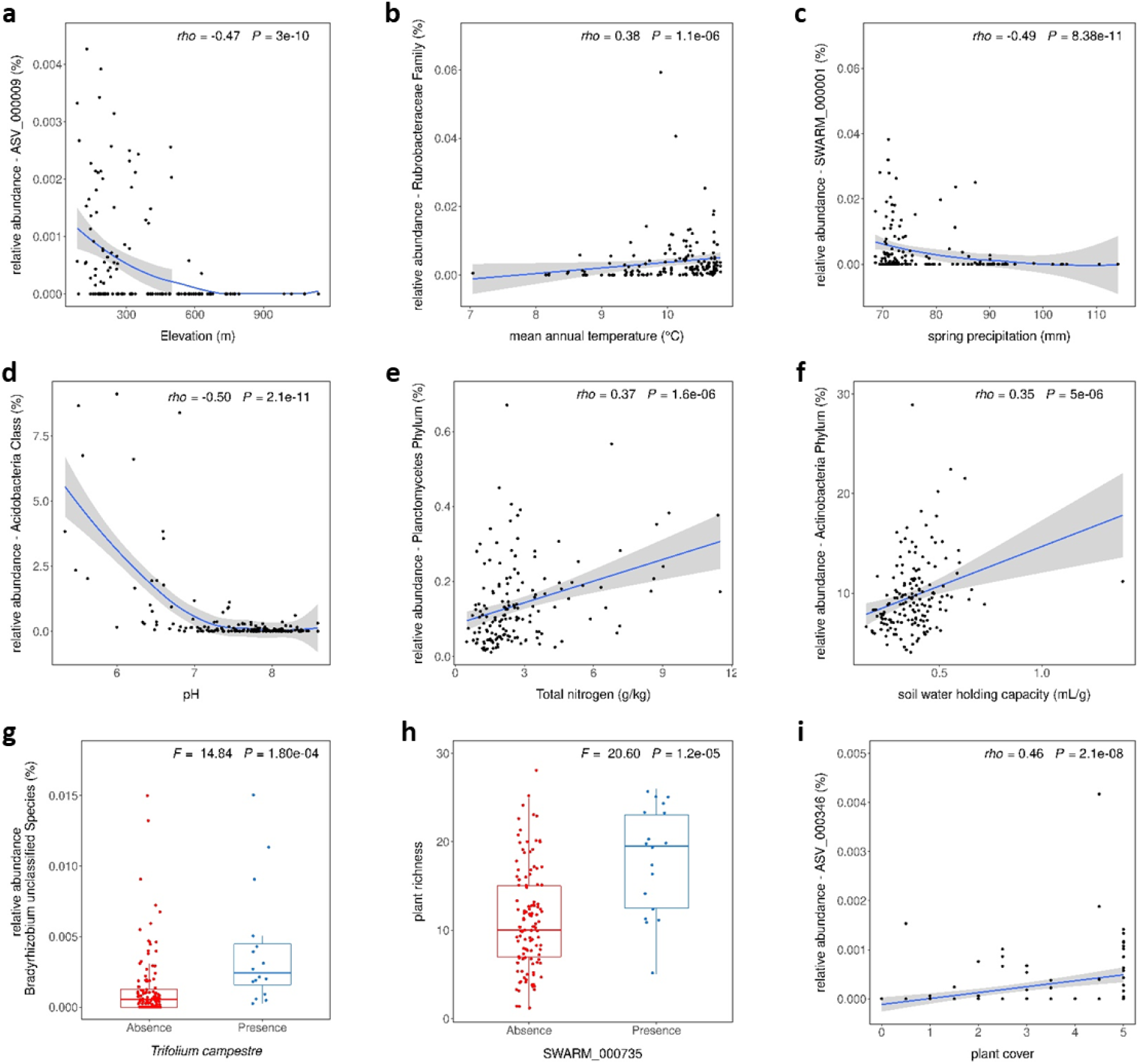
Illustration of relationships between soil taxonomic groups and non-microbial ecological variables. **a** Negative relationship between the relative abundance of ASV_000009 belonging to the family *Sphingomonadaceae* and elevation. **b** Positive relationship between the relative abundance of the family *Rubrobacteraceae* and mean annual temperature. **c** Negative relationship between the relative abundance of ASV_000001 belonging to the genus *Nitrospira* and spring precipitation. **d** Negative relationship between the relative abundance of the class *Acidobacteria* and pH. **e** Positive relationship between the relative abundance of the phylum *Planctomycetes* and soil total nitrogen content. **f** Positive relationship between the relative abundance of the phylum *Actinobacteria* and soil water holding capacity. **g** Positive relationship between the relative abundance of an OTU corresponding to a *Bradyrhizobium* species and the presence of the plant species *Trifolium campestre*. **h** Positive relationship between the presence of SWARM_000735 belonging to the genus *Variovorax* and the richness of plant communities. **i** Positive relationship between the relative abundance of ASV_000346 belonging to the order *Rhizobiales* and cover of plant communities.

## DISCUSSION

In this study, we aimed to estimate the relative effects of abiotic and biotic factors in explaining the diversity and composition of soil bacterial communities at diverse taxonomic levels. To address this objective, we used the highly taxonomically resolving *gyrB* gene instead of markers with a lower taxonomic resolution, such as the 16S rRNA gene [47, 49]. Interestingly, despite using two different markers to characterize soil bacterial communities, the proportion of sampling sites where the major phyla were present as well as their relative abundance, were similar at two geographical scales, i.e. the French scale based on the sequencing of a 16S amplicon [36] and the regional scale of the southwest of France based on the sequencing of a *gyrB* amplicon in this study. The characterization of soil bacterial communities down to the infraspecific level was combined with the characterization of a large and unique set of non-microbial variables, including (i) micronutrients that have only been recently considered in explaining variation in soil microbiota structure [44–46], and (ii) descriptors of plant communities (i.e. diversity, composition, and presence/absence of plant species) beyond the classification of plant communities in large categories of vegetation types [6].

### The relative effects of abiotic and biotic factors in explaining the structure of soil bacterial communities depend on the taxonomic level

In this study performed at a regional scale, soil bacterial communities at the lowest taxonomic levels appear mainly structured by abiotic factors that have very often been identified at larger geographical scales. These key abiotic factors include pH [24, 36, 61], temperature [16, 41], and precipitations [62–65]. Because temperature and precipitations influence biogeochemical processing occurring in the soil matrix [66, 67], climate change is expected to strongly affect ecosystem services provided by soil microbes that will, in turn, affect the performance of aboveground biotic communities [68–70]. In agreement with recent studies [44–46], soil bacterial composition micronutrients were also often associated with soil bacterial composition in the southwest of France. Notably, soil exchangeable manganese concentration might be a key abiotic factor in structuring soil bacterial communities. Interestingly, soil exchangeable manganese concentration was also identified as strongly associated with (i) the leaf and root bacterial microbiota of *A. thaliana* sampled in the natural sites investigated in this study [50], (ii) the soil fungal communities of natural sites sampled across Europe and inhabited by *A. thaliana* [71], and (iii) the bacterial, fungal and protist communities of maize field soils in eastern China [44]. In addition, exchangeable manganese has recently been demonstrated to strongly regulate carbon storage in the humus layer of boreal forests [72]. Because manganese is essential for plant performance [73], this micronutrient deserves further investigation to dissect its functional role in soil-plant-microbiota interactions.

At the highest taxonomic levels, soil bacterial communities appeared mainly structured by descriptors of plant communities. The positive relationship between soil bacterial diversity and plant diversity and cover has been reported in several natural ecosystems and agro-ecosystems and could result from a range of interactions including the exchange of carbon and nutrients but also the effects of bioaggressors [74–77]. The positive associations between soil bacterial species or infraspecies taxa and the presence of particular plant species are reminiscent of the host specificity documented for many plant pathogens [78] including bacterial and fungal species [79–83]. Host specificity has also been reported for commensal endophytic plant bacteria [84–88], albeit it remains controversial [89]. Root exudates are the interface of chemical communication and reciprocal benefits between plants and soil microbes [90, 91]. Therefore, plant host specificity of soil bacterial members may result from root exudate profiles that largely differ among plant species [92–95]. Interestingly, root exudate profiles exhibit plant phylogenetic signals and depend on edaphic conditions [92, 96]. This edaphic-dependent functional redundancy of root exudate profiles might explain that linear combinations of soil physicochemical properties and descriptors of plant communities better explain the presence of specific soil bacterial species than when considering soil physicochemical properties and descriptors of plant communities individually. In line with this result, the effect of crop diversity on soil microbiota composition was experimentally demonstrated to depend on abiotic factors, such as fertilization and soil moisture [97].

### The diversity of associations between soil microbiota and non-microbial variables

Although soil microbiota sampling was performed three years after the characterization of non-microbial factors, we still detected strong associations between individual microbial taxonomic groups and non-microbial factors. This might be explained by the stability of the soil bacterial microbiota of our 160 natural sites over a few years, as previously observed for soil samples collected over three successive years across 17 European sites inhabited by *A. thaliana* [71].

Many associations between soil microbiota and non-microbial variables identified in this study (Fig. 7) are supported by previous studies. For instance, the negative relationship between the relative abundance of ASV_000001 belonging to the genus *Nitrospira* and spring precipitation is in agreement with an increase in the relative abundance of the phylum *Nitrospirae* in conditions of decreased precipitation [98–100]. The positive relationship between the relative abundance of the family *Rubrobacteraceae* and mean annual temperature is consistent with members of *Rubrobacterales* and *Rubrobacteridae* being more abundant in extremely hot and dry locations, such as deserts and other arid regions [62, 101]. For soil physico-chemical properties, the negative relationship between the relative abundance of the class *Acidobacteria* and pH was also observed along a 1,500 m elevational gradient in the Taibai Mountain [38]. In addition, while the positive relationship between the relative abundance of the phylum *Planctomycetes* and soil total nitrogen content is consistent with *Planctomycetes* playing a major role in the global carbon and nitrogen cycle [102–104], the positive relationship between the relative abundance of the phylum *Actinobacteria* and soil water holding capacity is in line with the use of *Actinobacteria* to reduce water repellency in hydrophobic soils [8]. Finally, for descriptors of plant communities, the positive relationship between the relative abundance of an OTU corresponding to a *Bradyrhizobium* species and the presence of the legume *Trifolium campestre* is consistent with the observation of the predominance of *Bradyrhizobium* in the nodules of several species of the genus *Trifolium* in China [105]. In addition, in the low Arctic tundra, approximately 30–40% of microbes correlated with vascular plant coverage were assigned to *Burkholderiales*, *Chthoniobacterales*, and *Rhizobiales* [106]. Accordingly, our study revealed a positive relationship between the relative abundance of ASV_000346 belonging to the order *Rhizobiales* and the cover of plant communities.

### Future direction

Although our study was based on the characterization of 160 natural sites for a unique set of non-microbial variables, we have to be cautious that this list might still represent a glimpse of a larger number of non-microbial variables that could be linked to the structure of bacterial communities, including (i) variables that have previously been described as structuring soil bacterial communities, such as soil structure (e.g. macro aggregates), redox status, and relative humidity [24, 107], and (ii) variables linked to elevation (e.g. CO2 concentration, solar UV radiation) that was negatively associated with hundreds of individual microbial taxonomic groups across the eight taxonomic levels.

Determining the abiotic and biotic factors that influence the diversity, composition, and assemblage of soil microbiota relies on correlation approaches that are seldom followed by functional validation. Such functional validation requires the creation of large microbial collections that can be built with community-based culture (CBC) approaches [108, 109], which are facilitated by the development of ever more powerful microfluidics-related methodologies [110, 111]. By adopting a CBC approach, we recently established a collection of 7,259 bacterial colonies from the leaf compartment of 152 natural populations of *A. thaliana* located in the southwest of France [87]. A non-negligible fraction of members of this bacterial collection has also been identified in our soil samples, thereby allowing consideration of functionally validating associations identified in this study. For instance, the availability of three whole-genome sequenced strains of the Plant Growth-Promoting Bacteria *Paraburkholderia fungorum* from our leaf bacterial collection [87, 88] could be used to investigate the strong association identified between the presence of this bacterial species in soils and the presence of the plant species *Festuca rubra* (Supplementary Data 7).

Finally, the description of the bacterial communities of the 160 natural sites would benefit from the characterization of other microbial communities, including fungi and oomycetes. This may in turn help (i) to test whether the relative effects of abiotic and biotic factors in explaining the structure of soil fungal and oomycete communities also depend on the taxonomic level, and (ii) to estimate the variation of interkingdom interactions among sites and identify the underlying ecological drivers.

## ACKNOWLEDGMENTS

This work was performed at the LIPME of the Laboratoire d’Excellence (LABEX) entitled TULIP (ANR-10-LABX-41).

## COMPETING INTERESTS

The authors declare no competing financial interests.

## DATA AVAILABILITY STATEMENT

The *gyrB* gene amplicon sequencing raw data were deposited in the NCBI SRA database under the accession number SRP519501.

## SUPPLEMENTARY INFORMATION

**Supplementary Figure S1.** Association between the Shannon index of soil bacterial communities and non-microbial factors.

**Supplementary Table S1.** Metrics for the number of taxonomic groups and the number of reads before and after applying a filter based on the prevalence of sites.

**Supplementary Table S2.** Spearman’s correlation coefficients (*rho*) between the Shannon index and each of the 73 non-microbial ecological variables.

## SUPPLEMENTARY DATA

**Supplementary Data 1.** Non-microbial ecological characterization of the 160 sites.

**Supplementary Data 2.** *gyrB* database.

**Supplementary Data 3.** Number of reads per sample (N = 756) for each of the 327,387 ASVs. ‘Site’ corresponds to the 480 samples of the 160 natural sites. ‘citizen-CG’ and ‘citizen-IS’ correspond to the 276 samples of the citizen project.

**Supplementary Data 4.** Taxonomic affiliation for the 241,184 ASVs and the 212,837 SWARMs.

**Supplementary Data 5.** PERMANOVA results on the association between microbiota composition and non-microbial ecological variables at each of the eight taxonomic levels.

**Supplementary Data 6.** List of the 2,207 taxonomic groups present in more than 15 sites.

**Supplementary Data 7.** Spearman’s pairwise correlation coefficients (*rho*) between individual soil microbial taxonomic groups and non-microbial ecological variables.

**Supplementary Data 8.** sPLSR results on the association between individual soil microbial taxonomic groups and linear combinations of non-microbial ecological variables.

## Notes

### Competing Interest Statement

The authors have declared no competing interest.

