## Supplementary Information for "The relative effects of abiotic and biotic factors in explaining the structure of soil bacterial communities at diverse taxonomic levels"

**This file includes:**

- Supplementary Figures 1 and 2
- Supplementary Tables 1

**Supplementary Figure S1.** Association between Shannon index of soil bacterial communities and non-microbial factors. **a** Positive relationship between the presence of the plant species *Valerianella locusta* and the Shannon index of soil bacterial communities. **b** Negative relationship between the Shannon index of soil bacterial communities and the soil potassium concentration.

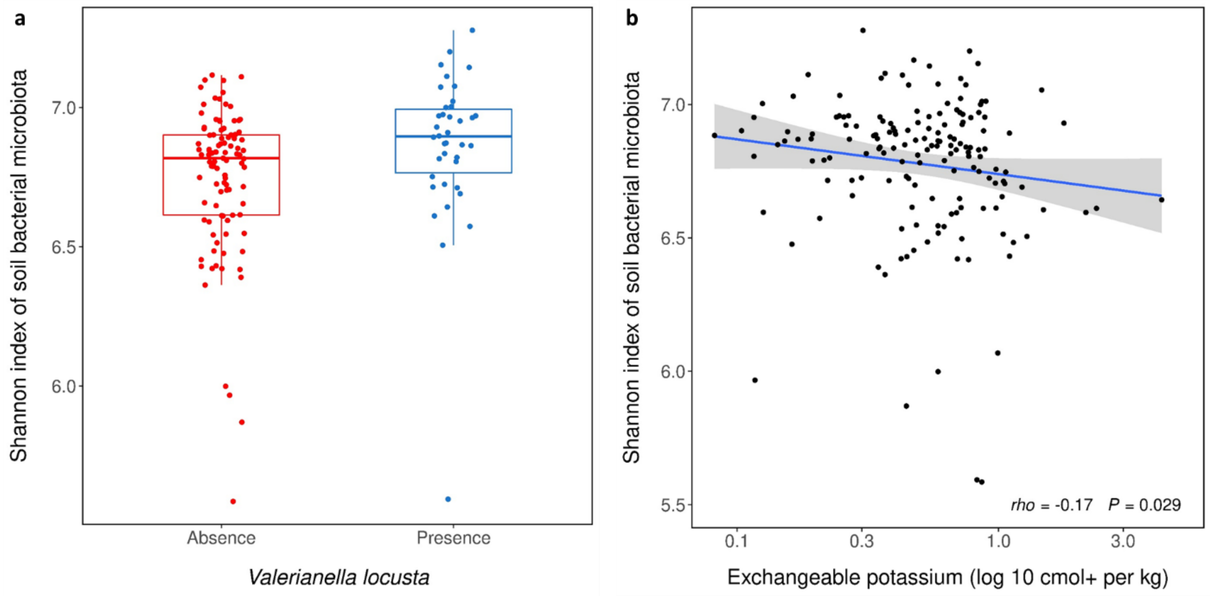

**Supplementary Table S1.** Spearman's correlation coefficients (*rho*) between the Shannon index and each of the 73 non-microbial ecological variables. See Supplementary Data 1 for a description of the 73 non-microbial ecological variables.

| non-microbial_variable | *rho* | *P* |
|---|---|---|
| Latitude | -0.15 | 0.06719 |
| Longitude | -0.09 | 0.26731 |
| Elevation | 0.03 | 0.72828 |
| MAT | 0.01 | 0.93110 |
| MCMT | 0.04 | 0.58113 |
| PPT_wt | -0.02 | 0.78396 |
| PPT_sm | -0.02 | 0.81384 |
| PPT_sp | -0.01 | 0.91844 |
| Nitrogen | -0.11 | 0.18378 |
| PPT_at | -0.01 | 0.85663 |
| CN | -0.02 | 0.77152 |
| pH | -0.06 | 0.47333 |
| Calcium | -0.14 | 0.08526 |
| Phosphore | -0.12 | 0.13651 |
| Magnesium | -0.11 | 0.16771 |
| Sodium | -0.12 | 0.12189 |
| Iron | -0.15 | 0.05263 |
| Potassium | -0.17 | 0.02930 |
| Aluminium | -0.07 | 0.36021 |
| WHC | 0.00 | 0.99215 |
| OC | -0.08 | 0.31818 |
| SOM | -0.08 | 0.31597 |
| Manganese | 0.09 | 0.26465 |
| plant_OTU1 | -0.02 | 0.79883 |
| plant_OTU3 | 0.03 | 0.72697 |
| plant_OTU4 | 0.01 | 0.87421 |
| plant_OTU7 | 0.14 | 0.11078 |
| plant_OTU8 | -0.08 | 0.37789 |
| plant_OTU10 | 0.03 | 0.73704 |
| plant_OTU15 | -0.02 | 0.77747 |
| plant_OTU16 | 0.10 | 0.24645 |
| plant_OTU18 | 0.15 | 0.08283 |
| plant_OTU20 | -0.03 | 0.74014 |
| plant_OTU27 | 0.09 | 0.31497 |
| plant_OTU46 | 0.25 | 0.00334 |
| plant_OTU49 | 0.01 | 0.88427 |
| plant_OTU65 | 0.13 | 0.12123 |

| non-microbial_variable | *rho* | *P* |
|---|---|---|
| plant_OTU67 | -0.05 | 0.57360 |
| plant_OTU71 | 0.07 | 0.44496 |
| plant_OTU72 | 0.02 | 0.84625 |
| plant_OTU78 | -0.08 | 0.34088 |
| plant_OTU83 | 0.10 | 0.22452 |
| plant_OTU87 | 0.04 | 0.63843 |
| plant_OTU100 | -0.14 | 0.10438 |
| plant_OTU88 | -0.01 | 0.92047 |
| plant_OTU109 | -0.06 | 0.45998 |
| plant_OTU113 | -0.04 | 0.61236 |
| plant_OTU114 | 0.01 | 0.87740 |
| plant_OTU132 | 0.16 | 0.06378 |
| plant_OTU136 | 0.03 | 0.76392 |
| plant_OTU143 | 0.10 | 0.24630 |
| plant_OTU145 | 0.09 | 0.31005 |
| plant_OTU146 | 0.13 | 0.12525 |
| plant_OTU147 | -0.04 | 0.65055 |
| plant_OTU149 | 0.02 | 0.77406 |
| plant_OTU154 | -0.12 | 0.14589 |
| plant_OTU159 | 0.06 | 0.50449 |
| plant_OTU179 | 0.09 | 0.31988 |
| plant_OTU192 | -0.18 | 0.03954 |
| plant_OTU196 | -0.11 | 0.18165 |
| plant_OTU198 | -0.10 | 0.25938 |
| plant_OTU202 | -0.02 | 0.79860 |
| plant_OTU203 | -0.08 | 0.38367 |
| plant_OTU204 | -0.18 | 0.03203 |
| plant_OTU216 | 0.04 | 0.66045 |
| plant_OTU223 | 0.16 | 0.06852 |
| plant_OTU234 | -0.02 | 0.80542 |
| plant_richness | 0.16 | 0.05479 |
| plant_Shannon | 0.19 | 0.02579 |
| plant_pcoa1 | 0.14 | 0.11384 |
| plant_pcoa2 | 0.09 | 0.31568 |
| plant_pcoa3 | 0.03 | 0.74681 |
| plant_cover | 0.20 | 0.01679 |

**Supplementary Table S2.** Metrics at each taxonomic level for the number of taxonomic groups and the number of reads before and after applying a filter based on the prevalence of sites.

| | Raw data | | After filtering (presence >15 sites) | |
|---|---|---|---|---|
| | **Number of groups** | **Number of Reads** | **Number of groups** | **Number of Reads** |
| **Phylum** | 27 | 3 503 024 | 18 | 3 502 510 |
| **Class** | 48 | 2 779 309 | 32 | 2 778 649 |
| **Order** | 82 | 1 572 894 | 50 | 1 571 147 |
| **Family** | 144 | 940 416 | 71 | 935 914 |
| **Genus** | 372 | 660 381 | 130 | 646 895 |
| **Species** | 563 | 605 313 | 127 | 583 490 |
| **SWARM** | 212837 | 6 778 650 | 1010 | 1 128 103 |
| **ASV** | 241184 | 6 778 650 | 769 | 824 198 |
